# Massive proteogenomic reanalysis of publicly available proteomic datasets of human tissues in search for protein recoding via adenosine-to-inosine RNA editing

**DOI:** 10.1101/2022.11.10.515815

**Authors:** Lev I. Levitsky, Mark V. Ivanov, Anton O. Goncharov, Anna A. Kliuchnikova, Julia A. Bubis, Anna A. Lobas, Elizaveta M. Solovyeva, Mikhail A. Pyatnitskiy, Ruslan K. Ovchinnikov, Mikhail S. Kukharsky, Tatiana E. Farafonova, Svetlana E. Novikova, Victor G. Zgoda, Irina A. Tarasova, Mikhail V. Gorshkov, Sergei A. Moshkovskii

**Affiliations:** V.L. Talrose Institute for Energy Problems of Chemical Physics, N.N. Semenov Federal Research Center for Chemical Physics, Russian Academy of Sciences, 38, bld. 2, Leninsky Prospect, 119334 Moscow, Russia; Federal Research and Clinical Center of Physical-Chemical Medicine, 1a, Malaya Pirogovskaya, 119435 Moscow, Russia; Pirogov Russian National Research Medical University, 1, Ostrovityanova, 117997 Moscow, Russia; Institute of Biomedical Chemistry, 10, Pogodinskaya, 119121 Moscow, Russia; Insilico Medicine Hong Kong Ltd., Hong Kong Science and Technology Park, New Territories, Hong Kong, China; Institute of Physiologically Active Compounds, Federal Research Center of Problems of Chemical Physics and Medicinal Chemistry, Russian Academy of Sciences, 1, Severny Proezd, Chernogolovka, 142432 Moscow Region, Russia

## Abstract

The proteogenomic search pipeline developed in this work has been applied for re-analysis of 40 publicly available shotgun proteomic datasets from various human tissues comprising more than 8,000 individual LC-MS/MS runs, of which 5442 .raw data files were processed in total. The scope of this re-analysis was focused on searching for ADAR-mediated RNA editing events, their clustering across samples of different origin, and classification. In total, 33 recoded protein sites were identified in 21 datasets. Of those, 18 sites were detected in at least two datasets representing the core human protein editome. In agreement with prior art works, neural and cancer tissues were found being enriched with recoded proteins. Quantitative analysis indicated that recoding of specific sites did not directly depend on the levels of ADAR enzymes or targeted proteins themselves, rather it was provided by differential and yet undescribed regulation of interaction of enzymes with mRNA. Nine recoding sites conservative between human and rodents were validated by targeted proteomics using stable isotope standards in murine brain cortex and cerebellum, and an additional one was validated in human cerebrospinal fluid. In addition to previous data of the same type from cancer proteomes, we provide a comprehensive catalog of recoding events caused by ADAR RNA editing in the human proteome.

## Introduction

The ability to obtain and access the entire set of multi-omic (genomic, transcriptomic, proteomic, metabolomic, interactomic, etc.) data in a fairly short time has revolutionized biological research in recent years and led to deeper understanding of the essential biological mechanisms of living organism functioning (Hasin, Seldin, and Lusis 2017). To model complex biological interactions, these systems biology studies rely on the integration of omics data using both experimental and computational methods (Akiyama 2021; Y. V. Sun and Hu 2016). The basic assumption behind this integration, or, most likely, the main, long-lasting hope, is the idea that it is the integration of the entire set of multidimensional data on a biological object that will make it possible to overcome the heterogeneity barriers of real biological systems, thus, reaching the tempting perspectives of preventive medicine and personalized therapeutic approaches to the treatment of essential human diseases (L. Hood 2013; L. Hood and Flores 2012; Alyass, Turcotte, and Meyre 2015; R. Chen and Snyder 2012, 2013; Labory et al. 2020). Nevertheless, while the possibilities of personalized medicine look like the prospect of a near future, the reality is still far from these hopes, primarily due to the limited capabilities to generate and analyze multidimensional data (Alyass, Turcotte, and Meyre 2015). Moreover, with the advent of the fifth generation of sequencers in genomics and high-performance high-resolution mass analyzers in proteomics in recent years, significant progress has been made in experimental data generation. As a result, the creation of computational resources and the analysis of data obtained continuously within the framework of global consortium projects is becoming the main obstacle to the development of systems biology and the transfer of its results to the practice of medical research (Auffray et al. 2016; Tenenbaum 2016). Generally speaking, generation of large-scale experimental omics data is quickly becoming a protocol-determined routine and not the Achilles heel of system biology (Shendure and Lieberman Aiden 2012; Shilo, Rossman, and Segal 2020).

In view of the above, one of the growing areas in system biology research are processing, analysis and interpretation of proteomic data and their integration with genome information in the context of hypothesis driven multi-omic biomarker discovery studies. Thus, developing computational approaches, applying statistical methods to process experimental proteomic data aiming at increasing the throughput of the analysis and solving the above-mentioned problem of false-positive identifications becomes one of the important if not crucial tasks in proteome bioinformatics. In the same context of personalized biomedicine, the problem of false positives is especially acute when searching for rare events of genomic mutations at the whole proteome level when the search space of possible candidates becomes enormously large (Nesvizhskii 2014).

At present, rapid development of new generation mass spectrometric technologies based on high-resolution mass analyzers has resulted in accumulation of significant amounts of experimental data of proteome-wide analyses submitted at the growing pace to the ProteomeXchange consortium (Deutsch et al. 2017). This makes it possible to obtain medically significant qualitative and quantitative information about polymorphisms and their alternative variants associated with diseases (cancer, diabetes, cardiovascular diseases) at the protein level. One of the most popular sources of proteomic data in ProteomeXchange is PRIDE (Perez-Riverol et al. 2019). The PRIDE database (PRoteomics IDEntifications database, https://www.ebi.ac.uk/pride) is a public repository of proteomic data obtained by mass spectrometry supported by the European Bioinformatics Institute (EMBL-EBI). Currently, it has well over 14,000 proteomic project datasets with half of them related to human samples (Deutsch et al. 2020). Such a large amount of data facilitates their re-use for applications beyond the originally intended ones (Perez-Riverol et al. 2015; Martens and Vizcaíno 2017). Proteogenomics, which combines multi-omic data sets, may be one of the growing focuses of these proteomic data re-use activities (Hari et al. 2022; Robin et al. 2018).

Adenosine-to-inosine RNA editing is one of the interesting areas of proteogenomic research and may involve data reanalysis to search for the associated protein sequence variants. This evolutionary ancient post-transcriptional modification of RNA is catalyzed by adenosine deaminases of ADAR family (Nigita, Veneziano, and Ferro 2015). ADAR isoforms are able to bind to double-stranded RNA (dsRNA) and deaminate adenosine residues which results in inosine formation. The latter nucleotide is more affine to cytosine, in contrast to the original adenosine paired with uridine. As a result, dsRNA structures are dislocated. In non-coding RNA parts, ADAR activity is responsible for inactivation of dsRNA, which can elicit innate immunity in the form of type I interferon response (A O Goncharov et al. 2019; George et al. 2011). In humans and rodents, ADAR1 isoform is thought to be responsible for this function. Loss-of-function mutations of this gene in humans cause a heavy hereditary disease, one of the forms of Aicardi-Goutieres syndrome, which is accompanied by autoimmune inflammation in different organs (Rice et al. 2012).

In parallel to the immunomodulatory function, ADARs can recode the protein sequences when they edit exonic RNA, changing adenosine to inosine which mimics guanosine in codons (Sommer et al. 1991). The proteoforms arising from RNA editing were deduced from RNA sequences soon after discovery of this modification type (Sommer et al. 1991). ADAR2 is an isoform encoded by a distinct gene in mammals, which is prone to editing exonic RNA sequences, thereby recoding proteins (Duan, Tang, and Lu 2022). Despite thousands of potential recoded sites being deduced from RNA sequencing, a few of them are characterized functionally. Thus, a gene knockout of ADAR2 which causes an early death by excitotoxicity in mice can be reversed by the genomic knock-in of a single Arg-607 residue into GRIA2 glutamate receptor subunit instead of glycine, thereby mimicking Gln-to-Arg substitution via RNA editing (Higuchi et al. 2000). In humans, ADAR2 mutations are, in most cases, fatal, most likely, due to deficiency in the same GRIA2 site substitution. Furthermore, a Q-to-R amino acid change in Ig repeat 22 of murine filamin A, a recoding site conservative between mammals, is shown to regulate smooth muscle contraction and also may modulate a health state of the animal (Jain et al. 2022).

In mammals, rare examples of functional protein recoding by RNA editing are described. It is suggested that many of potential recoding events represent just the side effects of dsRNA inactivation by ADAR and, thus, are functionally nonadaptive (Xu and Zhang 2014). In one of the first efforts to reveal the proteome-wide consequences of RNA editing, a few hundred of edited sites in ganglia and axons of the longfin inshore squid, the cephalopod characterized by extremely frequent A-to-I RNA editing of transcripts, were identified using shotgun proteomics approach (Liscovitch-Brauer et al. 2017). This was followed recently by identification of recoded proteins in the fruit fly brain (Kuznetsova et al. 2018; Kliuchnikova et al. 2020), as well as murine and human brains (Levitsky et al. 2019). While close to seventy recoded sites were identified in proteins out of less than a thousand sites predicted from RNA sequencing in the insects (Kuznetsova et al. 2018), this ratio was much lower in mammalian brains, with twenty to forty identified sites out of one to two thousand predicted ones, respectively (Levitsky et al. 2019). About a dozen of mammalian sites were conservative between species, which confirms that most of theoretically possible recoded proteins are nonadaptive and never reside in brain tissues for a long time.

Similarly, a recent re-analysis of proteomic data of the CPTAC cancer project yielded as few as seven reliably identified A-to-I RNA editing sites in a huge breast cancer dataset (Peng et al. 2018). Of those, a recoded alpha coatomer protein (COPA) was shown to be functionally involved in cancers (Peng et al. 2018; Jain et al. 2018). Interestingly, the same site was recently detected among six recoded tryptic peptides in a deep proteome analysis of murine liver (Wu et al. 2014).

In this work we performed re-analysis of a large cohort of publicly available proteomic datasets obtained for human samples of different origin from the PRIDE repository focusing on searching the ADAR-mediated RNA editing events.

## Methods

### Datasets

Datasets for analysis were selected manually from the PRIDE archive (Martens et al. 2005) based on a number of criteria. The selection was aimed at obtaining high-quality data acquired in data-dependent acquisition (DDA) mode representing a range of different organs/tissues in humans. In order to have a more uniform collection of data, we opted to work with datasets obtained using Orbitrap mass spectrometers. We focused on three major categories of samples to search for recoding events: cancer samples (glioma, gastric cancer, non-small-cell lung carcinoma (NSCLC), and laryngeal squamous cell carcinoma (LSCC), high-grade serous ovarian cancer (HGSOC), primary bladder urothelial carcinoma, triple-negative breast cancer (TNBC), diffuse large B-cell lymphoma (DLBCL), acute lymphoblastic leukemia, renal cell carcinoma, meningioma), neural tissue/fluid samples (cerebrospinal fluid (CSF) and brain tissue samples) and other tissues (muscle tissue, kidney, testis, umbilical cord mesenchymal stem cells, platelets, plasma and urine). All selected datasets are listed and characterized in Table S1 (Wingo et al. 2017; Carlyle et al. 2017; Macron et al. 2018; Guldbrandsen et al. 2014; Lan et al. 2018; Geyer et al. 2016; J. Sun et al. 2018; Hingst et al. 2018; Zellner et al. 2018; Clemente et al. 2018; Lee et al. 2018; Adav, Wei, et al. 2018; C. Y. Chen et al. 2018; Lietzén et al. 2018; Jin et al. 2018; Klaeger et al. 2017; Lawrence et al. 2015; Sabrkhany et al. 2018; Sathe et al. 2019; Adav, Subbaiaih, et al. 2018; Nassa et al. 2018; Zhao et al. 2016; Bereczki et al. 2018; Lepper et al. 2018; Worzfeld et al. 2018; Park et al. 2017; Murgia et al. 2017; Casanova et al. 2017; Cominetti et al. 2018; Oller Moreno et al. 2018; Sim et al. 2019; Papaioannou et al. 2019; Tran et al. 2018; Fel et al. 2019; Koch et al. 2019; Brummer et al. 2019; Fornecker et al. 2019; Lichti et al. 2015; Yang et al. 2019).

#### Annotation

Pre-selected datasets were annotated for further analysis and selection. For each RAW file, the following metadata were extracted: sample characteristics (subject, tissue, condition), experimental conditions (sample preparation, protease, depletion/enrichment methods, labeling method, fraction and replicate number) and instrumentation (separation method, mass spectrometer model, fragmentation method, mass analyzer used for MS/MS spectrum acquisition). A table was created for each dataset, with a row for each RAW file, capturing these metadata. The tables were filled based on the information provided in the corresponding publications. These tables were then used in the automated pipelines we built for data processing. All annotations are provided in Table S2 in Supporting Information.

The annotations were used in the following ways: (1) sample characteristics and experimental conditions (fractions, replicates) were used for grouping of files during postsearch processing; (2) experimental details were used for selection of appropriate search parameters (labeling and sample preparation methods affected the modification settings, mass spectrometer and MS2 mass analyzer affected precursor and fragment ion mass tolerances); (3) sample characteristics (disease, tissue type) were used for grouping of datasets in representing the final results (see Table 2 below, showing recoded isoforms specific to certain tissues identified across multiple datasets). All metadata were used for final selection of datasets for reanalysis.

**Table 2.**
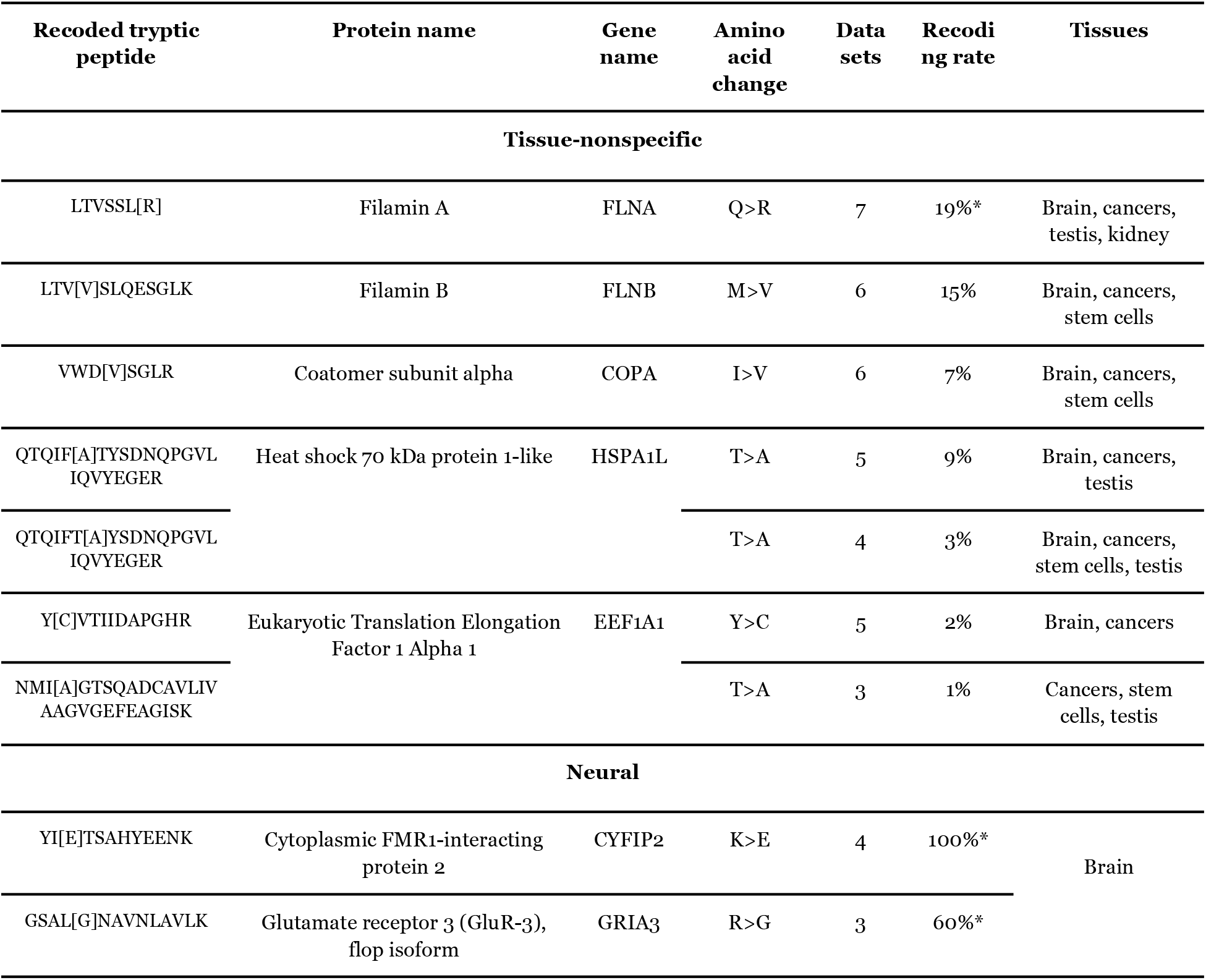
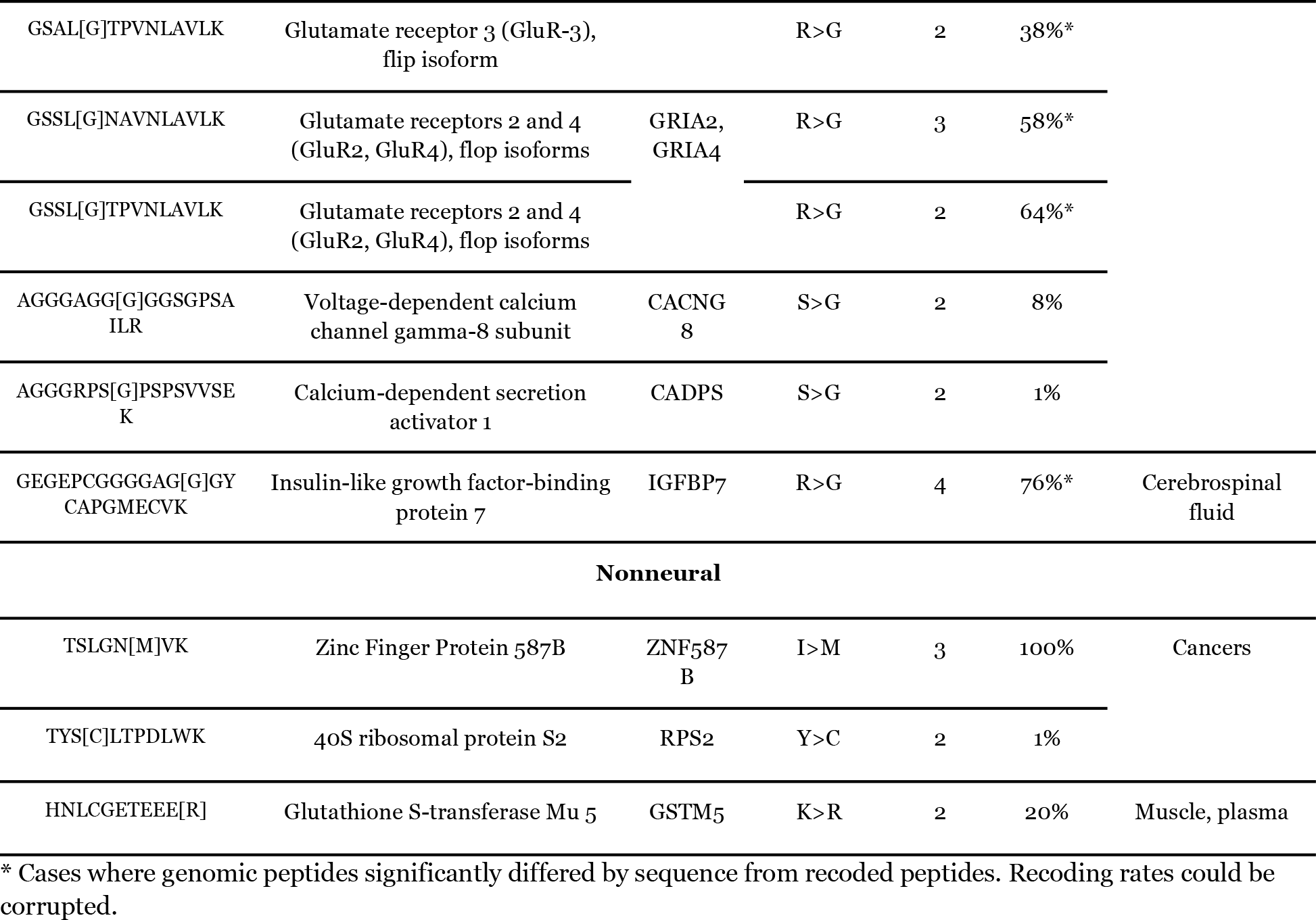
Protein sites recoded by ADAR RNA editing identified by proteogenomic analysis of human proteome datasets. Recoded sites were found in 21 of 40 datasets analyzed. Only those sites were represented which were identified at least in two datasets.Recoding ratio was calculated as a sum of PSMs of a recoded peptide divided by a sum of PSMs of the corresponding genome-encoded peptide. Substituted amino acid residues are outlined by square brackets. Additional information on datasets with recoded sites identified is provided in Table S8. In more detail, all identified recoded sites are characterized in Table S7.

### Proteogenomic pipeline

#### Сomposition of the database for search including sites of protein recoding via RNA editing

Two types of searches were conducted as part of the data processing pipeline, called wild-type and variant searches. SwissProt human protein database (20241 protein sequences) was used for wild-type searches. Decoy sequences were generated using the standard reverse approach and appended to the database. Variant searches were done using a combined database in peptide form rather than in protein form. Wild-type proteins were cleaved into tryptic peptides with up to two missed cleavages, while keeping the peptides of length 6 and above.

Database for protein recoding via RNA editing was also prepared in a peptide form based on the similar database as described above (Levitsky et al. 2019). Human genomic coordinates of RNA editing sites were obtained as described in (Levitsky et al. 2019). Additionally we introduced genetic variants with maximum allele frequency across populations exceeding one percent according to the gnomAD resource (Karczewski et al. 2020). The proteogenomic database also included results of potential ribosome stalling, i.e. peptides longer than six amino acids and ending with an N-terminal residue relative to the recoded residue. Final FASTA file contained a total of 17,874 recoded peptides. Decoy peptides were generated by reversing peptide sequences while keeping fixed N-term and C-term amino acids.

The Python implementation of the search pipeline is available at https://github.com/markmipt/bigdata_workflow under Apache 2.0 license. As input, the script needs a directory with raw files and preprocessed FASTA databases, and as output, it produces tables with FDR-controlled variant peptides, as well as intermediate files. The script contains nine consecutive steps as follows:

1. Conversion. Raw data are converted to standard mzML format using MSconvert from ProteoWizard pipeline (Kessner et al. 2008). Conversion is done using multiple filters: “peakPicking true 1-”, “MS2Deisotope”, “zeroSamples removeExtra” and “threshold absolute 1 most-intense”.
2. Wild search. All mzML files are searched with IdentiPy (Levitsky et al. 2018) search engine against the wild-type protein database. The parameters for the search engine are: 10 ppm precursor mass accuracy, 1 missed cleavage, carbamidomethylation of cysteine as fixed modification, minimal peptide length of six amino acids and autotune option. Fragment mass accuracy is either 0.03 or 0.3 Da depending on the type of mass analyzer used for measuring the fragmentation (MS2) spectra for a particular dataset. TMT mass tag at lysine and N-term of peptide is also set as a fixed modification for the corresponding datasets. IdentiPy search engine is run along with Dinosaur feature detection algorithm (Teleman et al. 2016) to extract MS1 peptide isotopic cluster intensities for identified MS2 spectra. Scavager post search algorithm (Ivanov et al. 2019) is used to process the output of IdentiPy.
3. Variant search. All the data are processed again using IdentiPy/Scavager combination to search against the peptide-based variant database.
4. Brute-force search. One hundred of top-scored wild-type proteins are chosen for every processed dataset file. These proteins and their decoy counterparts are saved as separate FASTA files (one for each run). Using these protein databases, the brute-force searches (Ivanov et al. 2018) are performed using IdentiPy search engine. Briefly, brute-force approach checks every possible single amino acid change for all wild-type tryptic peptides. The idea is that LC-MS/MS data is enriched with modified peptides from top scored proteins. We assume that the most identified “single amino acid substitutions’’ for these proteins are just artifacts produced by abundant modifications.
5. Variant tables generation. A table with variant peptides is generated for each file in the dataset. Variant peptides are sequences which belong only to variant proteins (target or decoy) and not shared with any wild proteins.
6. Group-specific FDR evaluation. All tables of variants are combined and the list of identifications is filtered to 5% group-specific FDR using posterior error probabilities calculated by Scavager.
7. Prosit MS/MS prediction. MS/MS peak intensities are predicted using PROSIT (Gessulat et al. 2019) for 5% FDR filtered variant peptides and for 1000 random wild-type peptides identified. Next, correlations between experimental and predicted MS/MS spectra are calculated and the threshold of correlation is calculated as the 5-th percentile for wild-type peptides. Variant peptides are filtered where correlations are below the threshold. The idea here is that modifications mimicking amino acid substitutions should result in lower correlations between predicted and experimental spectra.
8. DeepLC RT prediction. Theoretical retention times are predicted using DeepLC (Bouwmeester et al. 2021) for 5% FDR filtered variant peptides and for all identified wild-type peptides. The RT prediction errors are calculated as the difference between predicted and experimental peptide retention times. The distribution of RT errors for wild-type peptides is fitted as a sum of normal and uniform distributions. The mean shift and standard deviation are estimated and used to filter out variant peptides with RT errors outside of the 3 standard deviation range. The idea here is that modifications mimicking amino acid substitutions should result in a higher difference between predicted and experimental RT.
9. Final table composition. The final variant peptide table is composed, containing all information about the identifications, including the number of identified PSMs, best MS/MS correlation, lowest RT error, amino acid substitution rate in brute-force search, etc.

#### Additional filtering of the output of the proteogenomics pipeline

Each recoded peptide from the mutated database was searched against the UniProtKB/Swiss-Prot human reference proteome (UP000005640). Additionally for each peptide we generated all possible combinations of E>Q and D>N replacements. Peptides including all their variants that were matched in the reference proteome were discarded. Peptide variants which did not have either K or R at N-terminus, but were found in the reference proteome, were discarded as potential results of nonspecific cleavage. The only peptide variants considered were those which were not contained in the reference proteome sequences, nor any of their E>Q/D>N variants. The procedure was intended to avoid false reporting of genome-encoded peptides as recoded and resembled the so-called MS-Blast method (Shevchenko et al. 2001).

Additionally, peptide mass spectra of interest were visualized using xiSPEC spectrum viewer (Kolbowski, Combe, and Rappsilber 2018). The mass spectra were subjected to manual examination. Coverage of corresponding amino acid substitution by MS/MS fragments additionally indicated validity of a peptide spectrum match.

### Animals

Two transgenic mouse lines characterized by the proteinopathy in its nervous system were used for quantitative analysis of recoded proteins. The first of them represented mice expressing truncated FUS 1–359 protein. The production of this protein which is highly prone to aggregation leads to neurodegeneration, severe motor phenotype and premature death of animals at average age of 4 months (Shelkovnikova et al. 2013). The second mouse line was transgenic for human tau protein with the P301S mutation as described elsewhere (Allen et al. 2002). These mice demonstrated neurological phenotype dominated by a severe paraparesis accompanied by accumulation of hyperphosphorylated tau in the nervous system. The average lifespan of these mice in our facility was about 9 months. For both lines, all transgenic animals used for the experiment were asymptomatic but already had morphologically developed neural pathology as expected (Allen et al. 2002; Shelkovnikova et al. 2013).

We also used a transgenic mouse line generated recently in our facility which is yet unpublished. The line is called NEAT1_1Tg and does not express an exogenic protein. These mice harbor short isoform of long noncoding RNA NEAT1 which plays an important role in some cellular processes, such as transcription regulation in context of neurodegenerative and psychiatric diseases (Hirose et al. 2014; An, Williams, and Shelkovnikova 2018). It was shown that loss of Neat1 affects alternative splicing of genes important for the CNS function and implicated in neurological diseases (Kukharsky et al. 2020). NEAT1_1Tg mice are viable and have no overt phenotypic abnormalities despite some compromised behavioral symptoms (data not published).

Congenic mouse strains, CD1 for FUS 1-359 and C57Bl/6J for Tau P301S and NEAT1_1Tg, respectively, served as a wild type control. Animals matched by age and sex were used for both experimental and control groups. At age of 3, 5 and 6 months for FUS 1-359, Tau P301S and NEAT1_1Tg mice, respectively, cerebral cortices and cerebella were collected (Table S4), immediately frozen on dry ice and stored at −80° C until analysis was performed. Genotyping was performed by PCR analysis of lysates from ear biopsies using the following primer pairs: 5’-TCTTTGTGCAAGGCCTGGGT-3’ and 5’-AGAGCAAGACCTCTGCAGAG-3’ for FUS 1-359; 5’-AAGACGGGACTGGAAGCGATGAC-3’ and 5’-GCGGCAGTGTGCAAATAGTCTAC-3’ for Tau P301S; 5’-GGGACAACATTGACCAACGC-3’ and 5’-ACCACGGTCCATGAAGCATT-3’ for NEAT1_1Tg. All animal experiments were carried out in accordance with the Rules of Good Laboratory Practice in Russian Federation (2016).

### Murine brain sample preparation

Dissected cortices of one brain hemisphere and cerebellums were used as murine brain samples. After isolation and rinsing with saline, 150 μL of lysis buffer containing 2% (w/v) SDS, 0.1 M DTT, and 0.1 M Tris-HCl was added to each sample. Thereafter, sample was homogenized by heating (99 °C for 5 minutes) and ultrasonic sonication by Bandelin Sonopuls HD2070 ultrasonic homogenizer (Bandelin Electronic, Berlin, Germany) at 50% amplitude using 15 sec pulses for 5 min.

Methanol-chloroform precipitation was performed to separate proteins from interfering compounds. After drying, the protein pellet was resuspended in 100 μL of a solution containing 0.1% (w/v) ProteaseMAX Surfactant (Promega, USA), 50 mM ammonium bicarbonate, and 10% (v/v) acetonitrile (ACN) and stirred for 60 min at 550 rpm at room temperature. In the case of incomplete dissolution of the protein pellet, the solution was sonicated with 50% amplitude using 15 sec pulses until complete dissolution.

Protein concentration was measured using the bicinchoninic acid assay. 500 mM dithiothreitol (DTT) in 50 mM triethylammonium bicarbonate (TEABC) buffer was added to the samples containing 200 μg of protein to the final DTT concentration of 10 mM followed by incubation for 20 min at 300 rpm at 56 °C. Thereafter, 500 mM iodoacetamide (IAM) in 50 mM TEABC were added to the sample to the final IAM concentration of 10 mM. The mixture was incubated in the darkness at room temperature for 30 min.

The total resultant protein content was digested with trypsin (Trypsin Gold, Promega, USA). The enzyme was added at the ratio of 1:40 (w/w) to the total protein content, and the mixture was incubated overnight at 37 °C. Enzymatic digestion was terminated by addition of formic acid (5% w/v). Then, the samples were stirred at 500 rpm for 30 min at 45 °C followed by centrifugation at 15,700 g for 10 min at 20 °C. The supernatant was then added to the filter unit (30 kDa, Millipore, USA) and centrifuged at 13,400 g for 20 min at 20 °C. After that, 100 μL of 50% formic acid was added to the filter unit and the sample was centrifuged at 13,400 g for 20 min at 20 °C.

The total resultant peptide content was cleaned-up with Sep-Pak C18 cartridges (Waters, USA). The final peptide concentration was measured using Peptide Assay (Thermo Fisher Scientific, USA) on a NanoDrop spectrophotometer (Thermo Fisher Scientific, USA). The sample was dried using a vacuum concentrator (Concentrator 5301, Eppendorf, Germany) at 30 °C. Dried peptides were stored at −80 °C until the LC-MS/MS analysis.

### Human cerebrospinal fluid sample preparation

CSF samples were kindly provided by Dr. Victoria Shender and Prof. Vadim Govorun from the biobank of the Center of Physico-Chemical Medicine (Moscow, Russia). These samples were withdrawn from patients with various neurological diseases and were described in more detail earlier (Ziganshin et al. 2016). The pool of twenty samples was formed and then was concentrated on Amicon Ultra 0.5 mL Centrifugal filters MWCO 3 kDa (Merck, Germany) according to manufacturer’s instructions. Protein concentration was measured using the bicinchoninic acid assay. 150 μg of total protein were subjected to one-step reduction and alkylation, in a 50 mM triethylammonium bicarbonate buffer (TEAB) (Sigma-Aldrich, St. Louis, MO, USA) (pH 8.5) containing 25 mM tris(2-carboxyethyl)phosphine (TCEP) (Thermo Fisher Scientific, Waltham, MA, USA) and 40 mM chloroacetamide (CAA) (Sigma-Aldrich, St. Louis, MO, USA) at 80 °C for 40 min. The reaction mixture was diluted with 100 μL of 50 mM TEAB (pH 8.5), and the trypsin solution containing 3 μg of trypsin was added (Promega, Fitchburg, WI, USA), followed by incubation overnight at 37 °C. Hydrolysis was stopped by adding formic acid (Sigma-Aldrich, St. Louis, MO, USA) to a final concentration of 5%. The obtained peptide mix was spiked with SIS in ratio 28.5 fmol/μg of total protein followed by alkaline fractionation as described by Vavilov et al (Vavilov et al. 2020). The peptides were separated using a gradient of mobile phase containing 80% acetonitrile, 15 mM ammonia acetate in HPLC grade water, pH 9.3. The collected fractions were pooled into five samples containing the peptides eluted with 5-15, 15-30, 30-40, 40-50, and 50-60% of organic solvent. The samples were dried using a vacuum concentrator (Concentrator 5301, Eppendorf, Germany) at 30 °C. Dried peptides were stored at −80 °C until the LC-MS/MS analysis. The sample preparation procedure has been repeated in three technical replicates.

### Synthesis of stable isotope-labeled peptide standard

Peptides were synthesized by solid phase method using amino acid derivatives with 9-fluorenylmethyloxycarbonyl (Fmoc) protected α-amino groups (Novabiochem). The procedure was performed as described elsewhere (Kuznetsova et al. 2018). Resin with attached stable isotope-labeled lysine (L-Lys (Boc) (^13^C_6_, 99%; ^15^N_2_, 99%) 2-Cl-Trt, Cambridge Isotope Laboratories) was used for synthesis of peptides of CADPS (AGGGRPSSPSPSVVSE***K***^h^ and AGGGRPSGPSPSVVSE***K***^h^), GRIA3 (NAVNLAVL***K***^h^, GSALRNAVNLAVL***K***^h^ and GSALGNAVNLAVL***K***^h^), GRIA2 and GRIA4 proteins (TPVNLAVL***K***^h^, GSSLRTPVNLAVL***K***^h^ and GSSLGTPVNLAVL***K***^h^), Cyfip2 (TSAHYEEN***K***^h^ and YIETSAHYEEN***K***^h^) and IGFBP7 (GEGEPCGGGGAGGGYCAPGMECV***K***^h^, ITVVDALHEIPV***K***^h^ and GYCAPGMECV***K***^h^). Resin with attached stable isotope-labeled arginine (L-Arg (Pbf) (^13^C_6_, 99%; ^15^N_4_, 99%) 2-Cl-Trt, Cambridge Isotope Laboratories) was used for synthesis of peptides of CADPS (VNEEMYIE***R***^h^ and VNGEMYIE***R***^h^), Copa (VWDISGL***R***^h^ and VWDVSGL***R***^h^) and one of the peptides of IGFBP7 (GEGEPCGGGGAG***R***^h^). In more details, the peptides are shown in Table 1. Further steps of synthesis were also performed as described earlier (C. A. Hood et al. 2008).

**Table 1.**
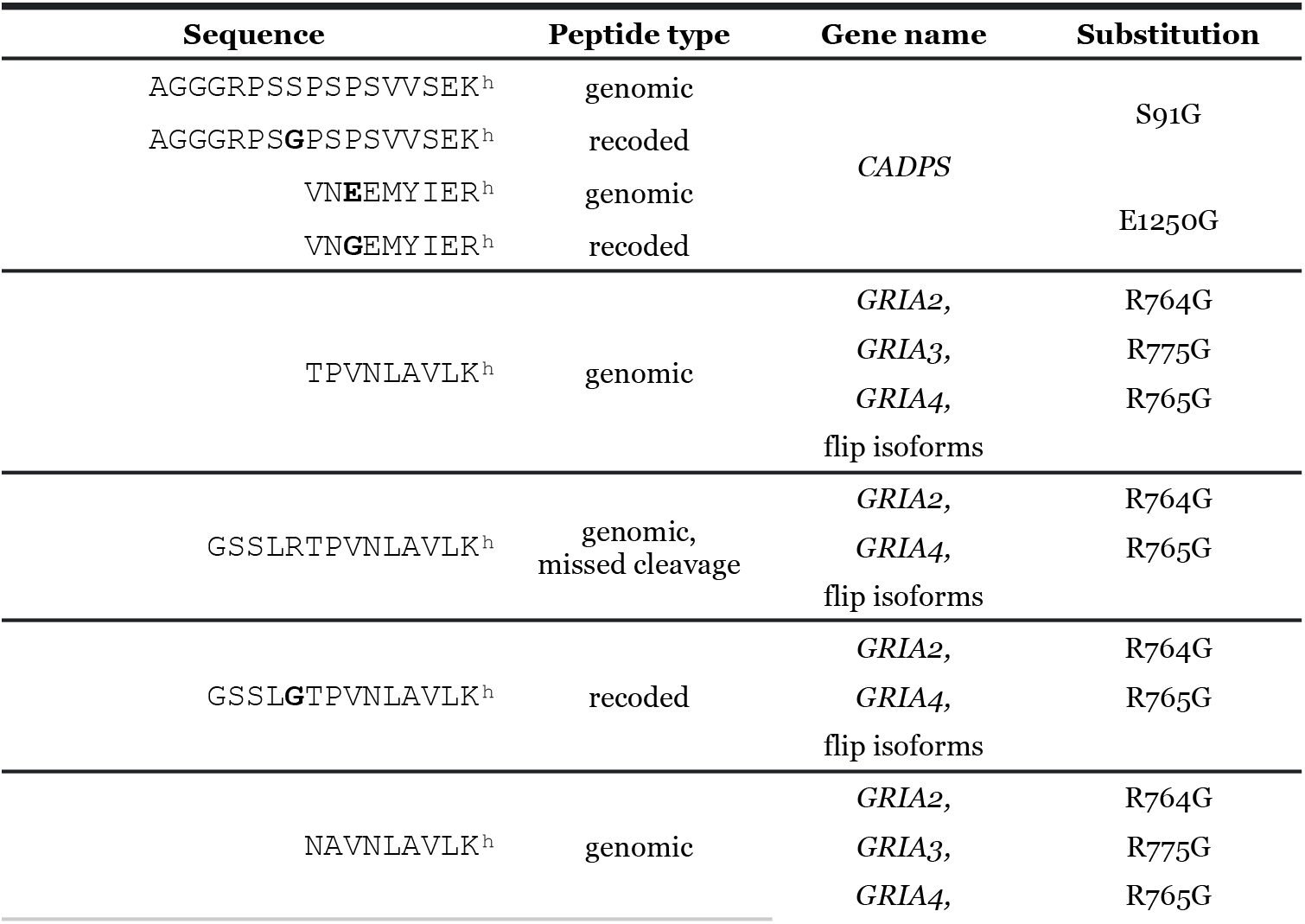
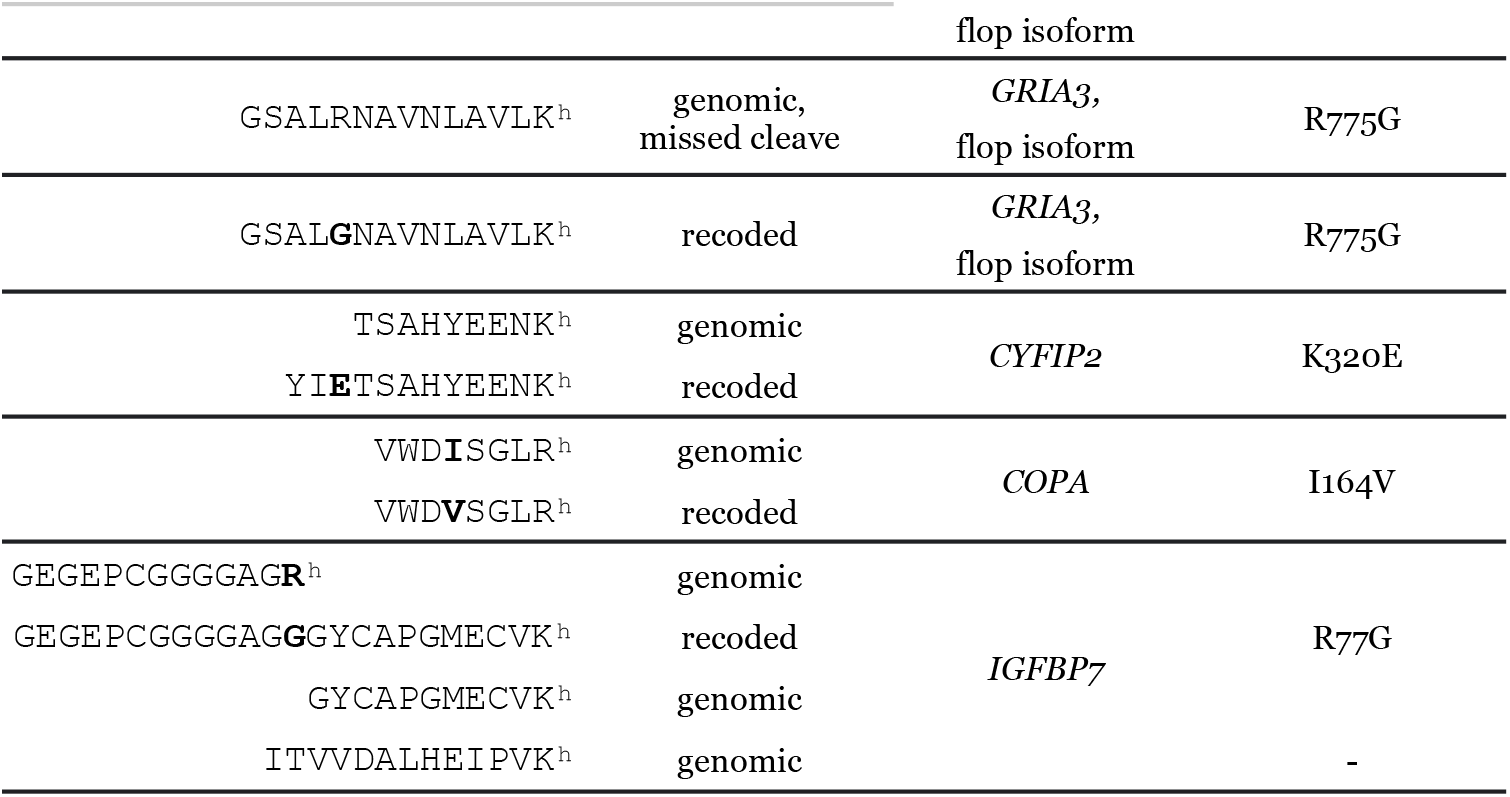
Tryptic peptides corresponding to genomically encoded and ADAR editing recoded protein sequences used for MRM-SIS analysis. These sites were chosen to be validated using multiple reaction monitoring with isotopically labeled standards (MRM-SIS). Amino acid numbering is represented for human genes, according to the first isoforms for each protein in the Uniprot database (UniProt Consortium 2021). The sites are identical and have the same numbers for mouse and human, with the exception of murine *Cadps* and *Gria3* genes, for which the corresponding amino acid substitutions are E1205G and R769G, respectively.

For quality control of the synthesis, LC-MS analysis was done using a chromatographic Agilent ChemStation 1200 series connected to an Agilent 1100 series LC/MSD Trap XCT Ultra mass spectrometer (Agilent, USA). Since some peptides contained methionines, the quality control also included manual inspection of the MS and MS/MS spectra for possible presence of the peaks produced by oxidized compounds. No such peaks were found in our case.

Absolute concentrations of synthesized peptides were determined using conventional amino acid analysis with their orthophthalic derivatives according to standard amino acid samples.

### MRM analysis

Murine and human samples were analyzed using a microflow Agilent 1200 HPLC system (Agilent Technologies, USA) connected to a triple quadrupole TSQ Quantiva (Thermo Scientific, USA) equipped with an electrospray ion source. Generally, 3 μL of each sample containing approximately 10 μg of total native peptides and 500 fmol of each stable isotope-labeled standard (SIS) peptide was loaded on an analytical column, Zorbax 300SB-C18 (5 μm, 150 × 0.3 mm) (Agilent Technologies, USA) and washed with 5% acetonitrile for 5 min at a flow rate of 20 μL/min. Peptides were separated using linear gradient from 95% solvent A (0.1% formic acid) and 5% solvent B (80% acetonitrile, 0.1% formic acid) to 40% solvent A and 60% solvent B over 30 minutes at a flow rate of 20 μL/minute.

The sets of transitions used for the analysis are shown in Table S5 for murine data and Table S6 for human samples. Capillary voltage was set at 4000 V, ion transfer tube temperature was 325 C°, vaporizer temperature was 40 °C, sheath and aux gas were set at 7 and 5 L/min, respectively. Isolation window was set to 0.7 Da for the first and the third quadrupole, and the cycle time was 1.2 s. MRM experiment was performed in a time-scheduled manner with a retention time window of 2 min for each precursor ion. Fragmentation of precursor ions was performed at 1.2 mTorr using collision energies calculated by Skyline 4.1 software (MacCoss Lab Software, USA) (Pino et al. 2020). Quantitative analysis of MRM data was also performed using Skyline 4.1 software. Quantification data were obtained from the “total ratio” numbers calculated by Skyline representing a weighted mean of the transition ratios, where the weight was the area of the internal standard. Up to five transitions were used for each peptide including the isotopically labeled standard peptide. Each MRM experiment was repeated in 3 technical replicates. The results were inspected using Skyline software to compare chromatographic profiles of endogenous peptide and stable-isotope labeled peptide. CV of transition intensity did not exceed 20% for technical runs.

## Results and discussion

### General overview of data and results

When processing a large collection of data from numerous sources, one of the main challenges is data heterogeneity. Because it is important to tailor search parameters to the data at hand, the data must be thoroughly annotated. Data annotation was done based on the referenced publications. For correct grouping of search results, the metadata had to contain sample characteristics: subject, tissue, condition, as well as the fraction and replicate number corresponding to each MS run. To determine the appropriate search parameters, experimental conditions must also be annotated. These include sample preparation methods, the protease used, depletion/enrichment methods, and labeling methods, all of which affect the modification settings. The chromatographic separation methods were annotated due to the fact that RT prediction is used in postsearch processing, and MS instrumentation characteristics (mass spectrometer model, fragmentation method, mass analyzer used for MS/MS spectrum acquisition) were annotated as well to ensure appropriate fragment ion type and m/z tolerance settings. All the above-mentioned characteristics were incorporated in SDRF-Proteomics – the PSI standard for sample metadata representation (Dai et al. 2021). The resulting file-level annotations are listed in Supplementary Table S2.

The search pipeline was developed with the aim of identifying variant peptides, while filtering out any identifications that can have alternative explanations. To this end, several protective measures were taken. At the database level, we included genetic variants with maximum allele frequency across populations exceeding one percent according to the gnomAD resource (Karczewski et al. 2020), as well as possible results of ribosome stalling. At the search level, a brute-force step was performed to identify frequent “substitutions’’ which could mimic chemical modifications or similar artifacts; all identified substitutions of these types were filtered out as unreliable. Additionally, all peptides which could be explained by E>Q or D>N substitutions were marked as unreliable, and peptide variants that could be explained by non-specific cleavage were designated a “grey zone”. Finally, at the post-search step, DeepLC and Prosit were employed for RT and MS/MS prediction, respectively. Both methods were used to filter out outliers. Finally, manual inspection of tandem mass spectra annotated with xiSPEC was performed.

Overall, the selected datasets contain more than 8,000 individual LC-MS/MS runs, of which 5442 .raw data files were processed to obtain the final results.

### Protein recoding via adenosine-to-inosine RNA editing in human proteomes

In twenty one of forty datasets searched, recoded sites were identified which met all selection criteria (Table S3). Not more than 18 peptides with recoded sites were found in each of them, containing up to several tens of thousands of genomic peptides, in correspondence to previous studies (Peng et al. 2018; Levitsky et al. 2019). In some of datasets, an insufficient depth of analysis was obviously a reason of undetected recoding, e.g., in body fluids such as ascite liquor, aqueous humor, blood plasma, and dermal interstitial fluid, as well as in datasets of hair shaft and T-cells (Table S3). Otherwise, some datasets with a lot of identified peptides also lack recoded identifications. In these tissues, a lower activity of ADAR enzymes, specifically ADAR2 which is responsible for the majority of protein sequence recoding (A O Goncharov et al. 2019). For example, liver, kidney, including urine cytology containing mostly renal epithelial cells, skeletal and cardiac muscle, and platelets represented tissues with low or absent protein recoding, which generally corresponded to the ADAR2 expression tissue profile from recent single cell transcriptomic study (Karlsson et al. 2021). Cancers were expectedly variable in terms of RNA editing-induced protein recoding. Eight of nine datasets originated from samples containing cancer cells had at least one recoded site identified. In contrast, a proteome of lung squamous cell carcinoma, being deep enough, nevertheless, did not contain detectable recoded sites.

In total, 33 recoded protein sites were identified in 21 datasets (Table S7). Of them, 18 sites were detected at least in two datasets, and, in turn, 20 datasets contained such re-identified sites. As an exception, a dataset of bronchoalveolar lavage was characterized by a single and unique recoded site in LRRFIP1 protein (Table S7). As an only observation of a recoded site may have no biological effect representing a sort of translational noise, further we focused on those eighteen sites which were found repeatedly (Table 2). All of them were reported on the protein levels in proteome-wide reports for different murine or human tissues (Peng et al. 2018; Levitsky et al. 2019).

Based on uniformed reanalysis of proteomes of various tissues performed here, we could classify repeated sites of recoding into three groups. The first of them was composed of the recoded sites widely distributed among different tissues. These sites included recoding of filamin A and B cytoskeletal proteins, coatomer α subunit (COPA), HSPA1L chaperone and translational elongation factor 1A1. Of them, at least two events are functionally characterized. Thus, Q/R editing of filamin A was responsible for regulation of the vascular contraction in murine models (Jain et al. 2018). Coatomer α I/V recoding was shown to destabilize the structure of the subunit which shortened the lifespan of this molecule. In absence of recoding, the more stable genomic type of coatomer α had oncogenic effects in hepatocellular carcinoma by activation of PI3K/AKT/mTOR pathway (Jain et al. 2018).

Another group of recoded sites consisted of events specific for neural tissues. These sites represented neural specific proteins, such as subunits of AMPA-type glutamate receptors (GRIA2, GRIA3, GRIA4), recognised as classical targets of A-to-I RNA editing (Sommer et al. 1991). The essential Q/R recoding site in the GRIA2 product occurred to be unfortunately unsuitable for tryptic digestion-based proteome analysis. Its recoding leads to formation of the tryptic digestion site, and resultant products of trypsinolysis are indistinguishable from semi-tryptic peptides or products of in-source fragmentation which makes this site a dark matter for this type of analysis. Instead, R/G recoding sites are perfectly identified in these subunits, both for *flip* and *flop* splice isoform (Table 2). This recoding event in GRIA2 was shown to accelerate the rate of channel opening and desensitization for heteromeric GRIA1/GRIA2 channel, where the latter was in the *flop* form (Wen, Lin, and Niu 2017). Of other recoded brain proteins, the voltage-dependent calcium channel gamma-8 subunit (CACNG8), or TARP γ8, should be mentioned, a protein functionally connected with AMPA glutamate receptor signalling. In the presence of TARP γ8, AMPA receptors change their electrophysiology and acquire resentisation by stabilizing conductive state (Carrillo et al. 2020). A functional connection between recoding of TARP γ8 and AMPA receptor subunits is not established yet, if any.

Cytoplasmic FMR1-interacting protein 2 (CYFIP2) is another protein recoded in the human brain. This protein is essential for normal brain development (Zhang et al. 2018), while its recoding by RNA editing is not characterized functionally. Further, CADPS is a synaptic protein also recoded in the brain. Interestingly, its fruit fly ortholog is also subject of recoding, although recoded sites in two homologs are not conservative (Kliuchnikova et al. 2020).

Separately from brain-specific recoded sites, an R/G site in insulin growth factor binding protein 7 (IGFBP7) was characteristic for cerebrospinal fluid as it was identified in all four datasets of this fluid involved (Table 2). Not much is known about the functionality of this site. Based on structural data in this protein, it was hypothesized that the R/G substitution in the insulin growth factor (IGF) binding domain could destabilize the interaction of the protein with its IGF ligand (Levanon et al. 2005). In addition, it was shown that R78G recoding along with K95R may modulate its susceptibility to proteolysis (Godfried Sie et al. 2012). Further, overexpression of ADAR2 was recently shown to facilitate lung fibrosis via connective tissue growth factor (CTGF) despite the exact mechanism of regulation of the latter by ADAR2 was not elucidated (Soundararajan et al. 2022). CTGF was supposed to be functionally connected with IGFBP7 and compete with the former for binding IGF isoforms (Wandji et al. 2000). Hypothetically, ADAR2 may influence CTGF by editing the IGFBP7 transcript and modulating interactions of its protein product. Because constant recoding of this site in the liquor seemed remarkable we chose to validate this site experimentally by targeted proteomic approach (see below).

In addition to some evolutionarily conserved sites in two groups described above, three sites with unknown function composed a group of recoding events identified outside neural tissues (Table 2).

Reanalysis of proteomic data in the context of protein recoding via RNA editing made it possible to estimate whether the extent of recoding is related to protein abundance. The first hypothesis is that in certain tissues with enough editing enzymes, such as ADAR2, transcripts are edited and proteins are recoded in proportion to their abundance. Otherwise, despite presence of the enzyme and the perceptive transcript, some trans-regulation can prevent its editing and recoding of the corresponding protein. In order to check these two options for different recoding sites, we clustered datasets and recoding sites based on relative recoding abundances, i.e. PSM counts of recoded peptides (Fig. 1A) and based on relative abundances of corresponding proteins, i.e. percentiles of each protein ranked by label-free quantitation values (Fig. 1B). Clustering also helped to analyze how tissues were distributed based on protein recoding profiles. Note that the PSM count for each recoded peptide is a very approximate metric which worked almost qualitatively in this clustering.

**Fig. 1.**
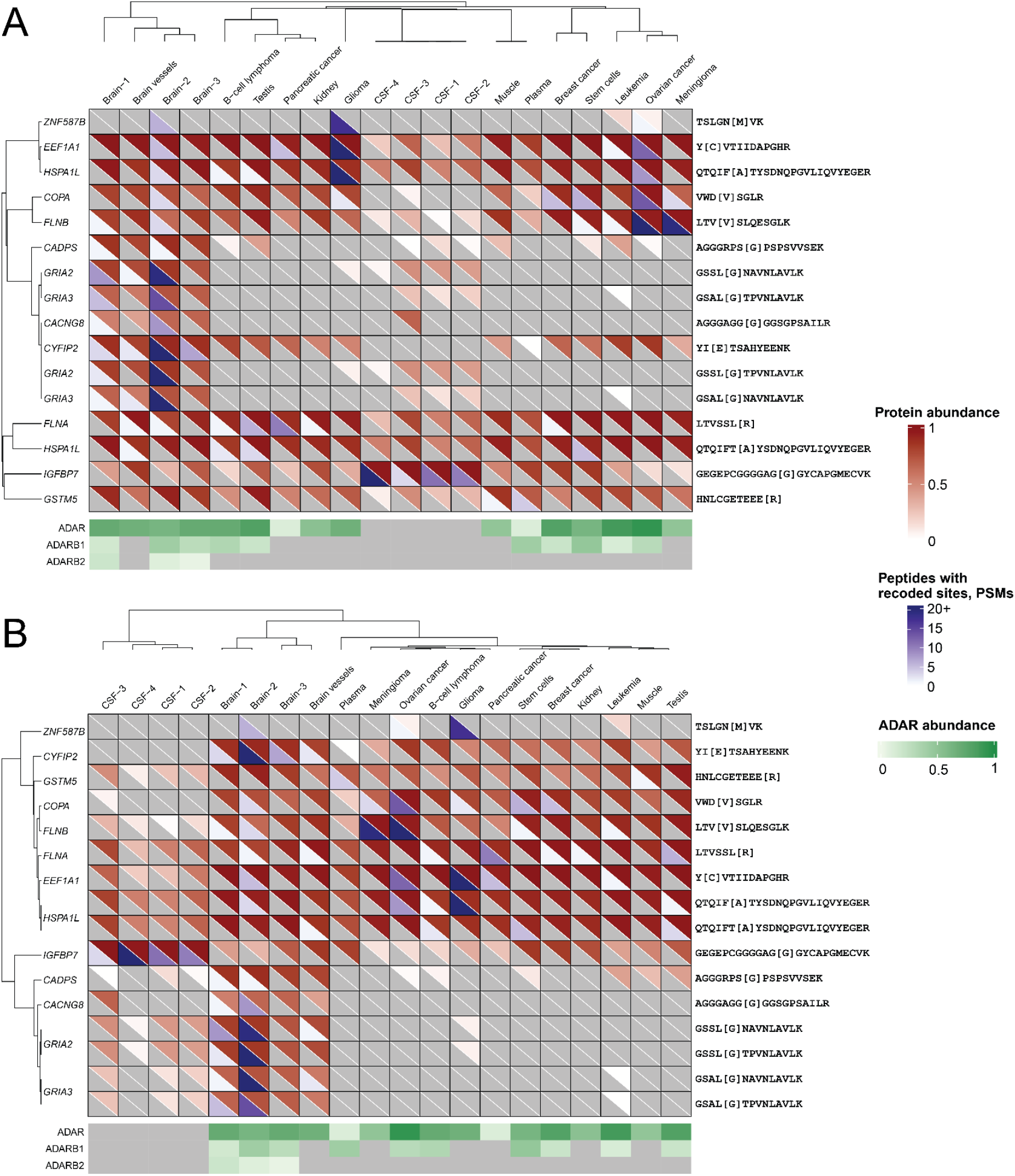
Heatmap of clustering of repeatedly identified ADAR-editing recoded sites and proteomic datasets where these sites were found. Datasets are designated by tissue of origin (for more detail, see Table S8). Recoded sites are designated by gene names of corresponding genes (for more detail, see Table 2). **A**: Heatmap of clustering based on total PSM numbers of each recoded site in each dataset. Exact PSM values are shown in Table S7. **B**: Heatmap of clustering based on label-free quantitation quantiles of each protein containing recoded site in each dataset. Exact quantile values are represented in Table S9. Label-free quantitation quantiles of three ADAR isoforms are shown in the bottom of each heatmap.

If the first of two scenario would happen and the recoding level would depend on ADAR activity and expression levels of edited products, both ways of clustering would lead to generally similar clusters. This similarity is observed on the heatmap, but with some important exclusions. Thus, a “neural” cluster based on recoding included recognized specific neural proteins such as ion channel subunits and CYFIP2 (Fig.1A). The neural cluster obtained based on protein abundance does not contain CYFIP2 which is produced in many other tissues (Fig.1B). However, its recoding is strictly neurospecific, which is presumably reached by regulation by some trans-acting factors on the level of mRNA. For example, SRSF9, a splicing modulator, was shown to interfere with ADAR2-dependent editing of some mRNAs by physical interaction with the enzyme (Shanmugam et al. 2018). Instead of CYFIP2, IGFBP7 joined other neurospecific protein in the abundance-based cluster, which could be explained by similarity of brain and cerebrospinal fluid (CSF) proteomes.

In addition, filamins A and B (FLNA and FLNB), which had very similar production profiles along the tissues and were co-clustered in Fig.1B, differed significantly in their recoding profiles. As in case of CYFIP2, it may be explained by the interplay between RNA splicing and editing (Anton O Goncharov et al. 2022).

In regard to tissue clustering based on protein recoding, a neural cluster is clearly distinct containing brain and brain vessel datasets (Fig.1A). Distinct recoding of filamins was a main differentiation factor between other two clusters containing cancer samples. Notably, a mesenchymal stem cell dataset was positioned tightly among cancer datasets. All CSF datasets are separated from others due to the presence of the specific IGFBP7 editing site.

### Validation and quantitation of evolutionary conserved recoded proteins in the murine brain

In our previous work, where recoded proteins were first identified in human and murine brains (Levitsky et al. 2019), we noticed that many of those sites and corresponding tryptic peptides identified in proteomes were identical between two species. Based on these fundings and many transcriptomic inventories (Hung et al. 2018), one can interpret omic data obtained with samples from both species interchangeably. Accordingly, the same kit of standards can be used for targeted proteomic analysis of these sites. Having murine models of neurodegenerative diseases in our access, we could pursue two tasks by quantifying recoded proteins in these models. First, existence of recoded sites on the proteome level could be validated. Most likely, the sites supported by murine brain data are expected to exist also in the human brain. Second, in three transgenic mouse lines expressing truncated *FUS* 1–359, human *tau* protein and truncated NEAT1 lncRNA, we could hypothesize some influence of the transgens to the ADAR editing and recoding. Jump ahead a bit, no effects of the transgenic manipulations were detected on levels of protein recoding which discarded the latter hypothesis. However, through checking the hypotheses, recoding events could be validated and quantified in proteomes of the brain cortex and the cerebellum.

In order to estimate the quantitative changes in recoding levels for the studied proteins, three groups of transgenic mice as above and their corresponding control groups were studied. Cerebellar and cortical samples were analyzed in three biological replicates for each sample. These brain tissues were characterized by high levels of ADAR RNA editing (Lo Giudice et al. 2020).

A recoding rate was calculated for the amino acid substitutions of interest (Table 2). This parameter was calculated for each sample from an individual animal, i.e. biological replicate based on averaged values of peptide levels for technical replicates. The recoding rate represented the recoded tryptic peptide concentration divided by a sum of the latter and the genomic peptide concentration. In case of multiple recoded isoforms known for glutamate receptor subunits, we tested one for each gene, specifically, the ‘flip’ isoform for GRIA2 and GRIA4 and the ‘flop’ isoform for GRIA3. Further, as in cases when genomically encoded tryptic peptides was shared between isoforms and even genes, the recoding rate was calculated in relation to the level of these peptide evenly divided between isoforms of interest. When one of the counterparts was not detected, the ratio may be conditionally defined as 0 or 1 (100%) in case of sole detection of genomic or recoded peptides, respectively. Expectedly, in these cases the dispersion of measurements cannot be estimated. The concentration of studied peptides in murine cortices and cerebelli is represented in Fig. S1.

Fig. 2 summarizes results on six recoded sites analyzed in mice. The recoding rate varied from 5-10% for *COPA* and *CADPS* sites up to 90% in R764/765G variant of glutamate receptor subunits *GRIA2* and *GRIA4* and supposed 100% in *CYFIP2* and *GRIA3* (cerebellum). As it was mentioned above, all three transgenic models provided remarkably similar results in terms of protein recoding. Thus, transgene manipulations with *FUS*, *tau* and *NEAT1* lncRNA did not influence ADAR RNA editing, at least, at the proteome level (Fig. 2).

**Figure 2.**
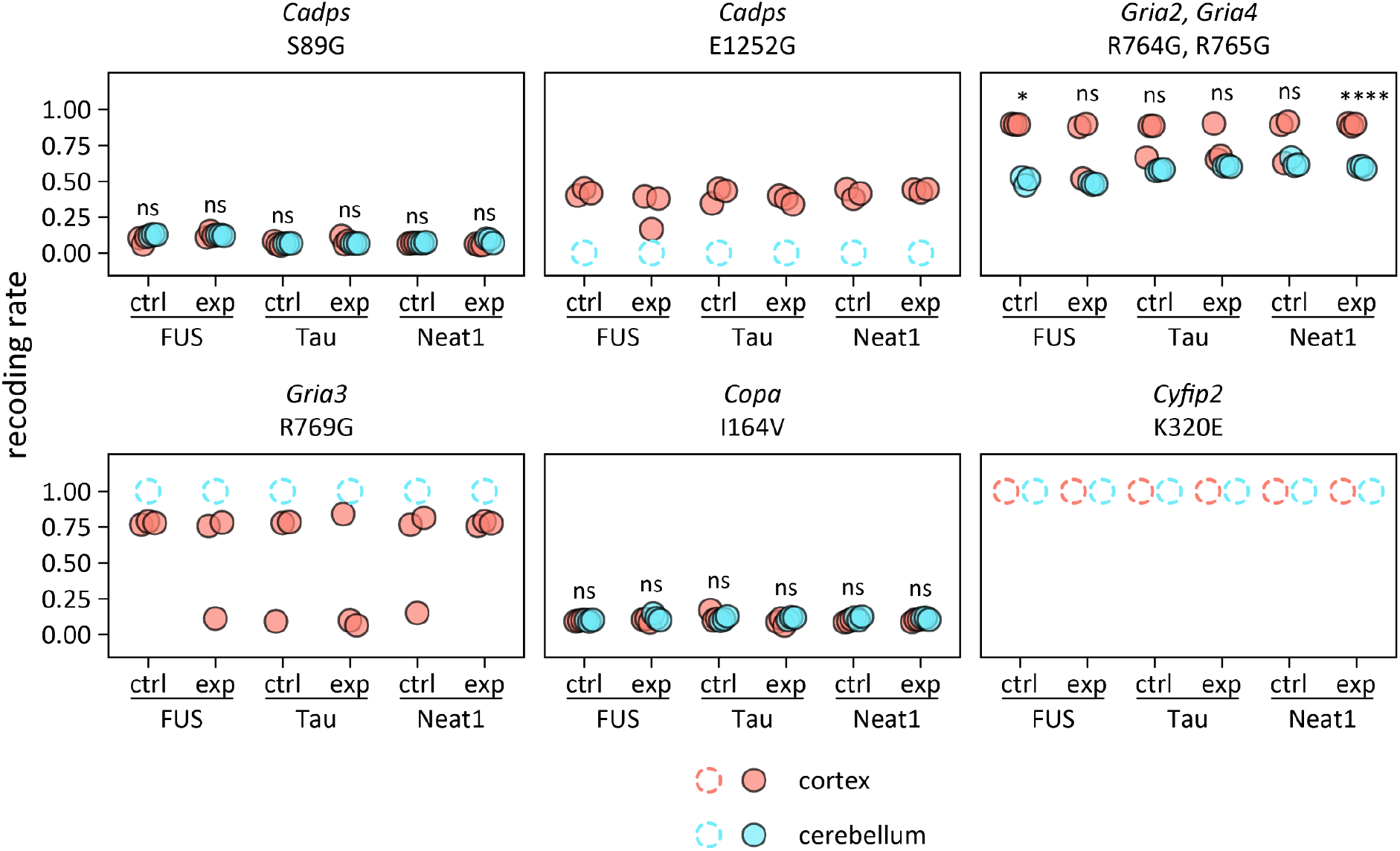
Recoding rates for sites of interest in cortex and cerebellum samples of FUS, *tau* and Neat1 transgenic mice and corresponding control strains. Transgenic interventions showed no statistically significant influence on recoding. A comparison between cortex and cerebellum outlined differences in R769G recoding in *Gria2* and *Gria4* products. For K320E recoding in the *Cyfip2* product a genomically encoded variant was not detected in both tissues, which most likely could be regarded as 100% editing. Similarly, for *Cadps* E1252G and *Gria3* R769G in cerebellum there was no detection of recoded and genomic variants, respectively. Hypothetical recoding rates estimated based on absent values are shown by dotted lines. Statistical significance was calculated using a *t*-test with Holm-Bonferroni adjustment for multiple comparisons. Asterisks indicate level of statistical significance: * – *p* ≤ 0.05, **** – *p* ≤ 0.001, ns – not significant, *p* > 0.05.

**Figure 3.**
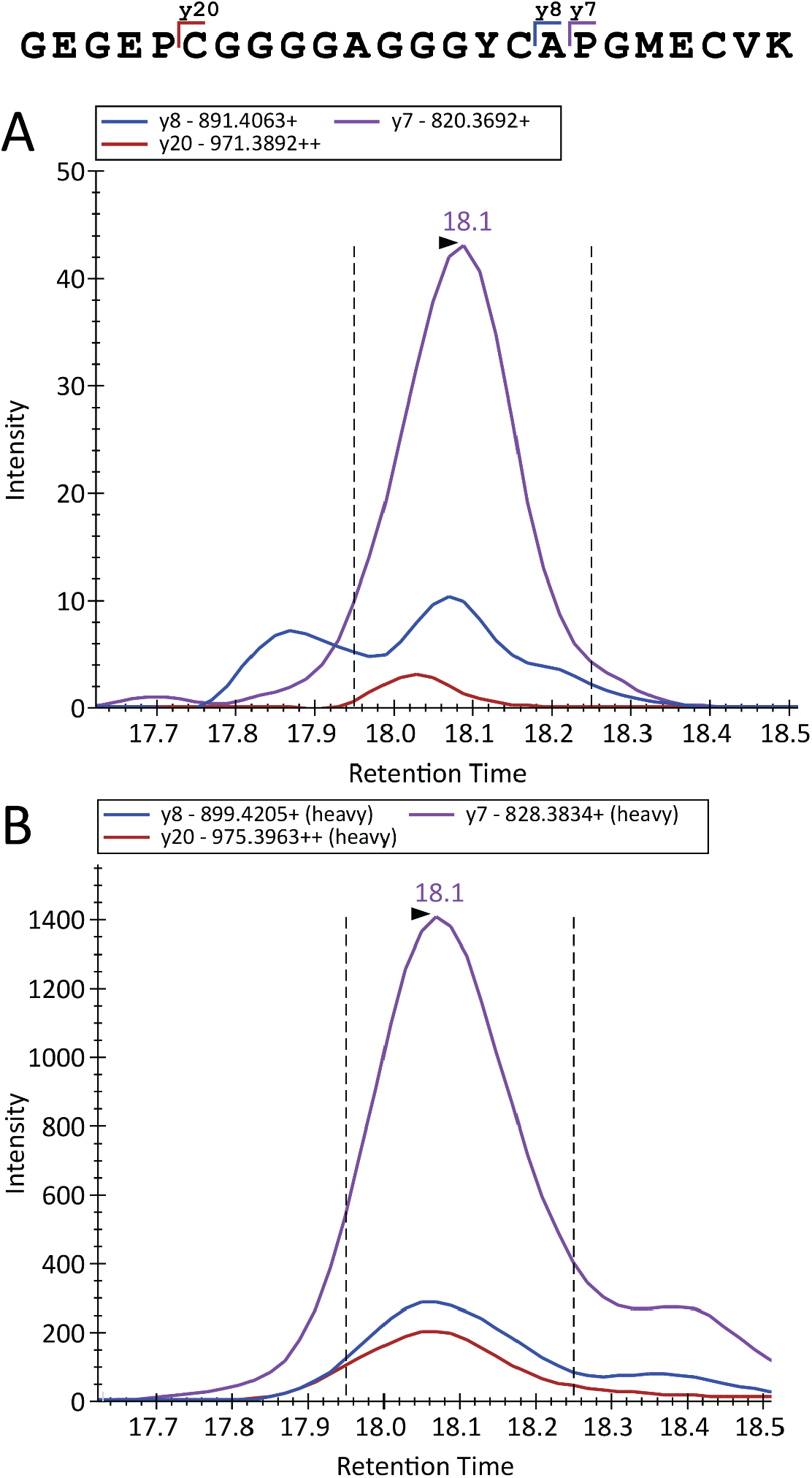
MRM chromatograms of the recoded peptide (GEGEPCGGGGAG[G]GYCAPGMECVK) of IGFPB7 protein detected in the trypsin-hydrolyzed human CSF pool. Data on the native peptide (A) and isotope-labeled peptide as internal standard (B). Spectra were visualized with Skyline 4.1 software. The retention time for the major peak is indicated by the arrow shown above the peak.

On the contrary, some differential recoding could be observed between cortex and cerebellum. For example, the recoding rate was higher in cortex for R764/765G variant of glutamate receptor subunits *GRIA2* and *GRIA4*. At the same time, in *GRIA3*, the homologous site was apparently fully recoded in cerebellum. Interestingly, two sites in the same gene, *CADPS*, behaved differently in cortex and cerebellum. The site S89E had the same recoding rate in both brain tissues, and E1252G was not recoded in cerebellum. Differential editing of two coding sites in the same transcript was recently observed in another gene, IGFBP7 (Godfried Sie et al. 2012). The site-specific regulation of ADAR editing in some transcripts was also described earlier, mostly being a result of interaction between RNA splicing and editing machineries (Anton O Goncharov et al. 2022).

### Validation of recoded Arg-78-Gly site of Insulin-like Growth Factor Binding Protein 7 in human cerebrospinal fluid

As it was shown before in this work and other studies (Godfried Sie et al. 2012; Levitsky et al. 2019; Macron et al. 2018), Insulin-like Growth Factor-Binding Protein 7 (IGFBP7) is extensively recoded specifically in CSF. In some rare cases, recoding of this product was also shown for other tissues, such as corpora striata of murine brains (Levitsky et al. 2019) and breast cancer (Peng et al. 2018). For validation of the only site of editing, R78G, detectable by conventional analysis of tryptic hydrolysates, targeted analysis of corresponding tryptic peptides was performed. As mentioned above, IGFPB7 is thought to participate in many biological processes including regulation of cell growth, apoptosis, and also involved in tumor suppression (Larsen et al. 2016). A peptide of interest containing the R78G recoding event with a sequence of GEGEPCGGGGAG[G]GYCAPGMECVK was aimed to analyze in pooled CSF samples, as well as two peptides corresponding to genomic sequence, GEGEPCGGGGAG[R] and GYCAPGMECVK, and, for additional control, the conservative peptide from other part of protein, ITVVDALHEIPVK.

As a result, ADAR-mediated recoding in IGFBP7 protein was confirmed, in spite of relatively low signal from the corresponding long peptide. Exemplary chromatograms of the targeted analysis are shown in Fig.3. In this case, the recoding rate could be calculated similarly as above, using averaged levels for two genomic peptides tested. Based on three technical replicates, the site R78G was characterized by the recoding rate of 0.61.

### Concluding remarks

Long ago, it was demonstrated for the first time how a single amino acid substitution in a single protein may result in a severe disease, a sickle cell anemia caused by D6V substitution in hemoglobin subunit β in that case (Ingram 1957). Just recently, a single conservative lysine-to-arginine substitution in neural TKTL1 was shown to affect propagation of neurons in neocortex which supposedly provided a superiority of Homo sapiens over Neanderthal in cognitive ability (Pinson et al. 2022).

RNA editing by RNA-dependent adenosine deaminases targets duplexes in various RNA molecules and destroys their regularity thereby impairing the reaction of innate immunity against these duplexes (George et al. 2016). At the same time, adenosine-to-inosine editing of mRNAs may recode resultant amino acid sequences which leads to translation of proteoforms with single amino acid polymorphisms. These proteoforms were shown to compete with genomically encoded counterparts and to have distinct functional properties (Jain et al. 2018).

In humans, in contrast to some invertebrates, a limited number of recoded proteins can be observed in the proteome. Only eighteen recoded sites were repeatedly identified in human tissue which is not a large number even taking into account limitations of shotgun proteomic methods. Based on tissue preferences they could be generally divided into non-specific and neuronal. A presence of recoded proteoforms at detectable levels advocates for their functional importance. Indeed, evidence accumulated on the impact of some of them on many physiological and cellular processes. The classical Q-to-R substitution in glutamate receptor subunits provides inhibition of neuronal excitability which is vital for the developing human brain (Tan et al. 2020). The recoded site in filamin A regulates contraction of vascular smooth muscles (Jain et al. 2022). A conservative change of isoleucine to valine in alpha coatomer provides potent tumor suppressor effects in liver cancer (Jain et al. 2018). In a similar fashion, other frequent recoded proteoforms, such as characteristic for the neuronal CYFIP2 protein, the IGFBP7 cytokine binder or the HSPA1L chaperone is supposed to participate in some pathways where they probably compete or otherwise interact with their genomic versions. Their mechanistic roles, if any, are still awaiting for elucidation.

Among prior art studies also using huge amounts of proteomic data to identify recoded proteins, a work by Peng et al analyzing cancer proteomes (Peng et al. 2018) and a recent study by Gabay et al focused on reliable analysis of ADAR RNA recoding in GTEx human samples (Gabay et al. 2022) should be mentioned. The latter work is concentrated on RNA sequencing data, but also reanalyzed proteomes in bulk manner, without tuning the search parameters for each dataset and without result filtration used here and by Peng et al. (Peng et al. 2018). Despite some differences in computational approaches, these works along with the present study form a comprehensive landscape of protein recoding via ADAR RNA editing.

## Institutional Review Board Statement

All subjects gave written informed consent in accordance with the Declaration of Helsinki. The human cerebrospinal fluid samples were used under a protocol approved by the Ethics Committees of corresponding clinical centers. For more details, see the previous paper which described the samples and their collection (Ziganshin et al. 2016).

## Informed Consent Statement

Informed consent was obtained from all subjects involved in the study.

## Data Availability Statement

All MRM mass spectrometry data are deposited to PASSEL (Farrah et al. 2012) (http://www.peptideatlas.org/passel/) under accession numbers PASS01741 for murine brain samples and PASS01759 for human cerebrospinal fluid, respectively.

## Acknowledgments

We would like to thank Prof. Vadim Govorun and Dr. Victoria Shender (Federal Research and Clinical Center of Physical-Chemical Medicine, Moscow, Russia) for providing human cerebrospinal fluid samples from the institutional biobank. Further, we thank the Center for Precision Genome Editing and Genetic Technologies for Biomedicine, Federal Research and Clinical Center of Physical-Chemical Medicine of the Federal Medical Biological Agency, which provided computational resources.

