## Supplemental Table 4 for "Massive proteogenomic reanalysis of publicly available proteomic datasets of human tissues in search for protein recoding via adenosine-to-inosine RNA editing"

**Table S4. Animals used for validation and quantitation of recoding events in the murine brain**

| **ID** | **Sex** | **Date of birthday, MM/DD/YYYY** | **Tissues were taken** | **Age** | **Father** | **Mother** | **Locus** |
| --- | --- | --- | --- | --- | --- | --- | --- |
| 101 | F | 07/07/2019 | 10/23/2019 | 108 | CD1 | FUS_SL_Homo | FUS_positive |
| 102 | F | 07/07/2019 | 10/23/2019 | 108 | CD1 | FUS_SL_Homo | FUS_positive |
| 103 | F | 07/07/2019 | 10/23/2019 | 108 | CD1 | FUS_SL_Homo | FUS_positive |
| 105 | F | 07/19/2019 | 10/23/2019 | 96 | CD1 | FUS_SL_Hemi | FUS_wt |
| 106 | F | 07/19/2019 | 10/23/2019 | 96 | CD1 | FUS_SL_Hemi | FUS_wt |
| 107 | F | 07/19/2019 | 10/23/2019 | 96 | CD1 | FUS_SL_Hemi | FUS_wt |
| 1 | M | 05/12/2019 | 10/13/2019 | 154 | Tau | Tau | Tau_positive |
| 2 | M | 05/12/2019 | 10/13/2019 | 154 | Tau | Tau | Tau_positive |
| 3 | M | 05/12/2019 | 10/13/2019 | 154 | Tau | Tau | Tau_positive |
| 5 | M | 05/12/2019 | 10/13/2019 | 154 | C57BL | C57BL | Tau_wt |
| 6 | M | 05/20/2019 | 10/13/2019 | 146 | C57BL | C57BL | Tau_wt |
| 7 | M | 05/20/2019 | 10/13/2019 | 146 | C57BL | C57BL | Tau_wt |
| 4781 | M | 04/10/2019 | 10/27/2019 | 200 | 5737 (5) | C57BL | Neat1_positive |
| 4785 | F | 04/10/2019 | 10/27/2019 | 200 | 5737 (5) | C57BL | Neat1_positive |
| 4786 | F | 04/10/2019 | 10/27/2019 | 200 | 5737 (5) | C57BL | Neat1_positive |
| 4825 | M | 04/10/2019 | 10/27/2019 | 200 | 5780 (1) | C57BL | Neat1_wt |
| 4829 | F | 04/10/2019 | 10/27/2019 | 200 | 5780 (1) | C57BL | Neat1_wt |
| 4830 | F | 04/10/2019 | 10/27/2019 | 200 | 5780 (1) | C57BL | Neat1_wt |
