## Supplementary figures and images for "Massive proteogenomic reanalysis of publicly available proteomic datasets of human tissues in search for protein recoding via adenosine-to-inosine RNA editing"

### Supplemental Figure S1

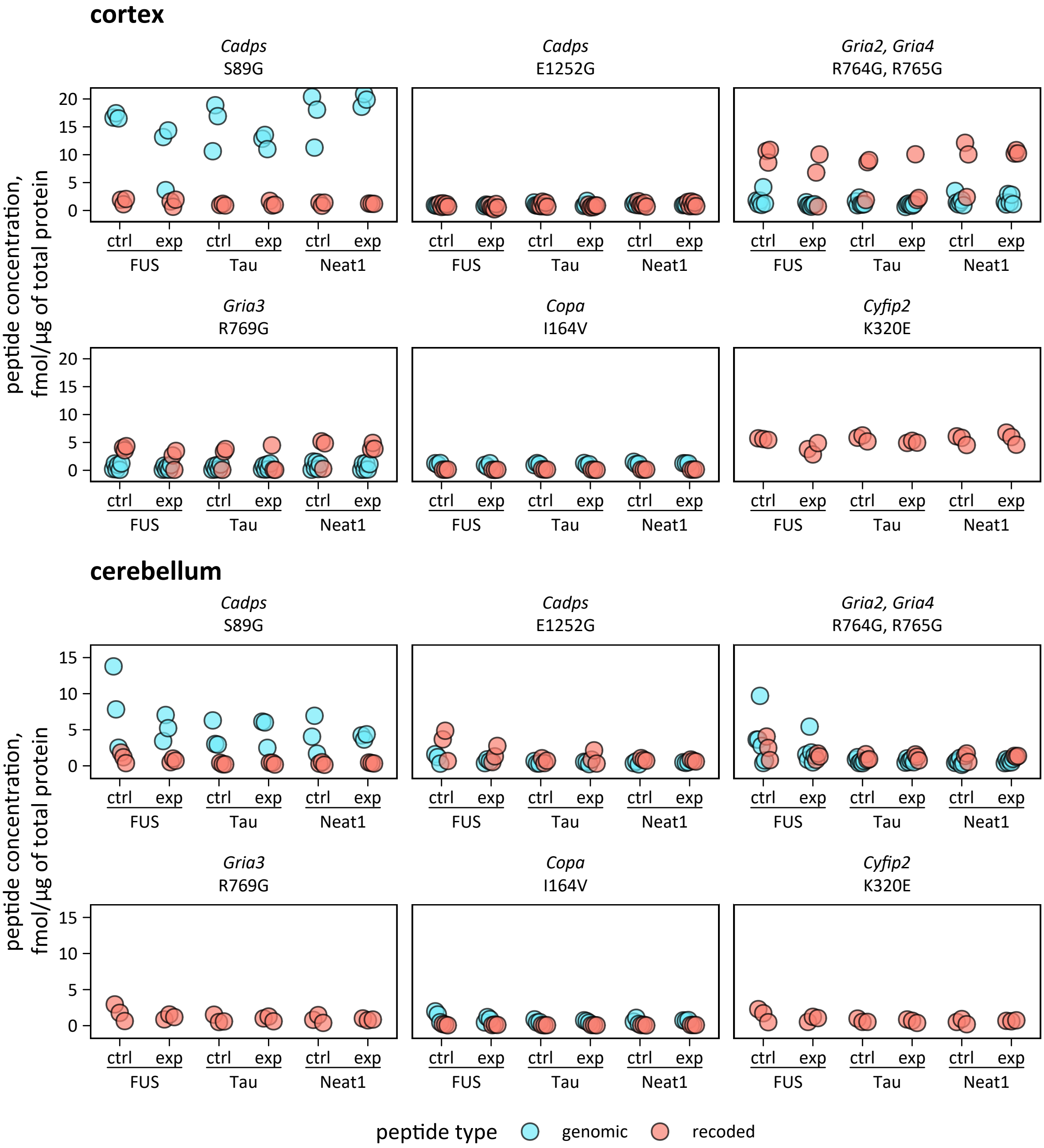
